# From minutes to bounds: A probabilistic UV-C control and a shape-only morphological fingerprint for postharvest *Colletotrichum*

**DOI:** 10.1101/2025.11.29.691312

**Authors:** Ezekiel Ahn, Insuck Baek, Seunghyun Lim, Amelia Lovelace, Minhyeok Cha, Moon S. Kim, Sunchung Park, Lyndel W. Meinhardt

## Abstract

Postharvest losses in high-value horticultural crops such as cacao and coffee are often driven by *Colletotrichum* spp. and other latent fruit pathogens. Ultraviolet-C (UV-C) is increasingly deployed as a chemical-free postharvest technology, yet prescriptions remain framed in minutes rather than in probabilistic guarantees of disease control and safety. We present a dual framework that (1) establishes conservative confidence bounds on survival and (2) validates a “shape-only” morphological fingerprint. This approach addresses the biological complexity of host-pathogen interactions by quantifying isolate-specific heterogeneity, rather than averaging it away. We utilized a large *Colletotrichum* dataset (n = 5,363) from cacao and coffee, spanning diverse treatments including UV-C, UV-B, and sonication. First, focusing on the Coffee UV-C cohort (∼10 min), we quantified this heterogeneity; the most conservative Clopper–Pearson upper 95% bound on survival reached 1.000, highlighting partial survival events (e.g., isolate P24-88) under otherwise high-efficacy conditions. This probabilistic framework captures the “tail-risk” of biological resilience instead of assuming complete kill. Second, we trained machine learning models on the full dataset using only geometric features (e.g., aspect ratio, asymmetry), explicitly excluding all primary size metrics. Serving as a rapid physiological indicator of UV-induced stress, these “shape-only” models successfully predicted pathogen host-origin (Accuracy ≈ 0.93) and post-treatment survival (R² ≈ 0.74). The signal’s ability to generalize across UV-B and sonication confirms that geometry, not just growth reduction, carries a robust and transferable physiological stress signature. This work provides a device-agnostic, probabilistic control platform, replacing time-based heuristics with quantitative guarantees and a generalizable, shape-based diagnostic.

**Highlights:**

- A conservative upper 95 % confidence bound (on survival = 1.000) was observed for the Coffee UV-C (∼10 min) cohort, revealing strong isolate-specific heterogeneity rather than a universal kill guarantee
- Isolate-specific heterogeneity (e.g., localized P24-88 survival) was quantified rather than averaged away.
- A size-free morphological fingerprint predicted host-origin (Accuracy ≈ 0.93) and survival (R² ≈ 0.74).
- The shape-only signal generalized across diverse stressors, including UV-C, UV-B, and sonication.
- A probabilistic, device-agnostic control framework replaces traditional time-based heuristics.

## 1. Introduction

Sanitation protocols for Ultraviolet-C (UV-C) in postharvest systems are widely adopted as a chemical-free control strategy, yet their prescriptions are often framed as “minutes” of exposure rather than as robust, probabilistic guarantees of efficacy (Ferreira et al., 2021). Practitioners require predictable margins of safety, but the operational outcomes of UV-C are confounded by a matrix of variables. Efficacy depends not only on total dose (fluence), but also on dose-rate (intensity), and is significantly modulated by fruit surface characteristics and packaging materials (Yu et al., 2024, 2023). Recent in vitro assessments have further highlighted that fungal inactivation kinetics differ remarkably between conidia and hyphal fragments, and between suspension and surface treatments (Bellino et al., 2025). This complexity leads to inconsistent results, where in vitro success fails to translate to in vivo application, and high doses risk damaging the commodity itself (Terao et al., 2015). This high variability, which manifests in our data as significant isolate-specific heterogeneity, necessitates a new framework that moves beyond simple time– or dose-based heuristics and embraces probabilistic, quantifiable guarantees.

In the context of postharvest supply chains, small changes in the efficacy of such treatments translate into large differences in food loss, especially for high-value horticultural commodities where latent *Colletotrichum* infections trigger anthracnose and stem-end rots during storage and transport. Conventional UV-C recipes are thus part of a broader portfolio of non-chemical preservation technologies that aim to reduce postharvest waste while maintaining marketable quality and food safety. However, the biological basis of these technologies, including isolate-specific tolerance, inoculum burden, and stress-induced morphological adaptation, remains under-quantified, limiting their scalability and transferability across devices and packhouses.

A second challenge lies in how fungal stress is measured. Traditional antifungal readouts often collapse the complex phenotypic response into a single, trivial metric: colony area (size). This overlooks the rich information encoded in fungal geometry. Abiotic stress, including UV radiation, is known to induce profound changes in fungal morphology at the cellular and colony level (Pietras et al., 2025). This link between stress response and morphological development is deeply rooted in fungal biology, with stress-response regulators directly governing morphological structures (Zhang et al., 2016). Consequently, the field of quantitative phenotyping is increasingly moving toward combining advanced imaging with artificial intelligence (AI) and machine learning (ML) to capture this complexity (Cembrowska-Lech et al., 2023). Recent studies have successfully applied deep learning and Random Forest algorithms to detect and classify fungal spores of Trichoderma and tomato pathogens with high accuracy (>95%), validating microscopic image processing as a reliable diagnostic tool (Javidan et al., 2024; Soltani Nezhad et al., 2024; Zhang et al., 2021). Recent studies now routinely use fungal “morphotypes” and geometric features, in combination with ML, to classify fungal isolates and quantify disease, confirming that morphology is a robust indicator of underlying biology (Gomez-Caro et al., 2022; Leiva et al., 2024; Upadhaya et al., 2025).

Here we present a unified platform that addresses both challenges. We utilized a large *Colletotrichum* dataset (n = 5,363) from cacao and coffee, intentionally spanning diverse treatments including UV-C, UV-B, and sonication. First, focusing specifically on the Coffee UV-C cohort (∼10 min), we quantify a conservative upper 95 % confidence bound on survival (= 1.000), revealing strong isolate-specific heterogeneity and the absence of a universal kill guarantee under high-inoculum, nutrient-rich conditions. Second, we trained ML models on the full dataset using only geometric features (e.g., aspect ratio, asymmetry), explicitly excluding all primary size metrics. These “shape-only” models successfully predicted pathogen host-origin (Accuracy ≈ 0.93) and post-treatment survival (R² ≈ 0.74). The signal’s ability to generalize across UV-B and sonication confirms that fungal geometry carries a robust and transferable stress signature. In addition to statistics, physics offers a device-agnostic bridge. By treating UV inactivation as a single-hit process and anchoring “minutes” to fluence and photon counts, one can invert confidence bounds on survival into lower-bound lethal cross-sections and hence risk-bounded times for target inactivation levels. To bridge the gap between biological variability and engineering control, this work replaces heuristic “minutes” with quantitative guarantees and elevates shape to a device-agnostic, generalizable diagnostic for fungal control.

## 2. Materials and methods

### 2.1. Fungal datasets and two-panel design

This study utilized a combined dataset (n = 5,363 observations) based on a two-panel design to test both generalizability and specific efficacy. The Cacao panel (n = 4,769), which served as our discovery cohort for fingerprint generalizability, was sourced from a comprehensive prior study (Baek et al., 2025b). This dataset included four isolates of *Colletotrichum gloeosporioides* (CGH17, CGH34, CGH38, CGH53) and one *Pestalotiopsis* sp. isolate (CGH5), originally obtained from the Cocoa Research Institute of Ghana, and subjected to diverse stressors (UV-C 275 nm, UV-B 305 nm, and sonication) as described therein.

The Coffee panel (n = 594), which served as our UV-C validation cohort, was generated specifically for the present study under strictly controlled conditions. For this, we utilized six *Colletotrichum* isolates (P23-55, P24-83, P24-85, P24-88, P24-192, and P24-193) originally sourced from *Coffee arabica* plants provided by Dr. Lisa M. Keith (USDA-ARS, Daniel K. Inouye U.S. Pacific Basin Agricultural Research Center) (Baek et al., 2025c).

### 2.2. Fungal inoculum preparation

Both panels followed the standardized inoculum preparation protocol detailed in Baek et al. (2025a). In brief, all isolates were maintained on potato dextrose agar (PDA) at 24°C in the dark and cultured for 10 days. Conidia were harvested by flooding plates with sterile distilled water (0.05% Tween 20) and filtering the suspension through sterile cheesecloth. The conidial concentration was determined using a hemocytometer and adjusted to 1 × 10^6^ conidia/mL. For the Cacao panel, 6–7 spots of 10 µL inoculum were placed onto each PDA plate (Baek et al., 2025b). For the newly tested Coffee panel, six spots of 10 µL inoculum were spot-inoculated onto PDA plates. All plates for this cohort were prepared, stored, and treated under an identical protocol.

### 2.3. UV stress application and dosimetry

The treatment modalities were applied differentially to the two panels. The sourced Cacao panel data reflected the diverse stressors and exposure times (e.g., UV-C 275 nm for 1-30 min; UV-B 305 nm for 5-30 min; sonication) required for the generalizability analysis (Baek et al., 2025b).

In contrast, the newly generated Coffee panel experiment was restricted to untreated controls and a single high-efficacy UV-C exposure (∼10 min, ∼600 s) designed to test our kill-guarantee hypothesis. All UV-C treatments for this cohort were performed using the custom-built, programmable UV exposure system previously described in Baek et al. (2025a). This system utilized top-mounted 275 nm UV-C LED modules and a computer-controlled moving stage. The intensity was monitored with a radiometer (Opsytec Dr. Gröbel GmbH) and adjusted to 0.58 mW/cm², ensuring that all replicates received uniform and identical dosimetry (Bolton and Linden, 2003).

### 2.4. Image analysis and morphometric phenotyping

Plates from both panels were imaged 48 hours post-inoculation (Baek et al., 2025b). Fungal colony morphology was characterized using SmartGrain software (v1.3) (Tanabata et al., 2012). We standardized seven key traits across both datasets: area (total pixel area), perimeter, length (longest axis), width (longest axis perpendicular to length), length-to-width ratio (LWR), circularity, and Euclidean distance between the axis intersection and the colony’s center of gravity (IS&CG).

### 2.5. Probabilistic confidence-bound framework

To move from “minutes” to “bounds,” we performed a probabilistic analysis focused specifically on the newly generated Coffee UV-C cohort. First, colony area was normalized to a survival ratio. This was computationally defined as Area_treated_/(mean(Area)_control_ + 10^−9^) to prevent division-by-zero errors if a control mean was zero. For binary classification (alive/not-alive), a stringent a priori threshold (τ) was set: a colony was defined as “alive” only if its survival ratio was ≥ 0.05 (5%). This conservative threshold prevents trace residues from being misclassified as survivors. We then computed the exact Clopper-Pearson 95% confidence interval (CI) for the true proportion of “alive” colonies within the ∼10 min window (Clopper and Pearson, 1934).

### 2.6. Multivariate and machine learning framework

All subsequent multivariate analyses were performed on the full dataset (n = 5,363) using Python (v3.11). Core libraries included pandas for data structuring and processing, statsmodels for ANOVA (Seabold and Perktold, 2010), SciPy for statistics (Virtanen et al., 2020), and Matplotlib for plotting (Hunter, 2007).

Principal Component Analysis (PCA): PCA was performed on log-transformed, standardized features to visualize the separation of pathosystems. This analysis focused only on the “shape-only” features (LWR, circularity, IS&CG) (Jolliffe and Cadima, 2016).

Machine Learning: To quantify the predictive power of the “shape-only fingerprint,” we trained two machine learning models (scikit-learn) (Pedregosa et al., 2011). To create this “size-free” fingerprint, we explicitly excluded all primary size metrics (area, length, width, perimeter) from the feature set, retaining only geometric features (LWR, circularity, IS&CG) and time/dose information. First, a Gradient Boosting Classifier (Friedman, 2001) was trained to distinguish host origin (Cacao vs. Coffee). Then, a Gradient Boosting Regressor (Friedman, 2001) was trained to predict the continuous survival ratio. For model validation, all models used plate-grouped cross-validation (Group K Fold) to prevent data leakage from repeated measurements on the same plate. Performance was reported using Accuracy and Macro-F1 (for imbalance) and R² / RMSE.

### 2.7. Physics-informed translation from bounds to times and energy

At 275 nm, the ∼10 min exposure used in the Coffee cohort was *I* = 0.58 mW cm^−2^ for t = 600 s, giving H = I ⋅ t = 348 mJ cm^−2^. Photon energy is E_hν_ = hc/λ and thus the photon fluence is N = H/E_hν_ = 4.82 × 10^17^ photons cm^−2^. We assume a single-hit inactivation process with survival S = exp(−σN). Using the isolate-wise Clopper–Pearson upper 95 % bound on survival at ∼10 min (S_up_), we obtain a conservative lower bound on the lethal cross-section: 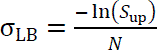

For a target inactivation fraction x ∈ {0.90, 0.99, 0.999} at irradiance I, the conservative time is 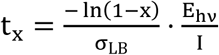

For practical planning with inoculum density ρ (conidia cm⁻²), the energy per expected kill at ∼10 min is upper-bounded by 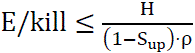 which highlights the burden penalty at higher ρ.

## 3. Results

### 3.1. Probabilistic confidence bounds and isolate-specific heterogeneity

Our primary practical objective was to move from “minutes” to “probabilistic bounds” for a common postharvest UV-C control scenario. We focused on the Coffee UV-C cohort (n = 594), which contained untreated controls and ∼10 min exposures. We defined a stringent binary criterion for survival (a normalized survival ratio ≥ 0.05) to conservatively classify any ambiguous colony growth as “alive”.

Applying exact Clopper-Pearson 95% CIs to the proportion of “alive” colonies in this ∼10 min window, we found the maximum upper 95% bound on survival reached 1.000 (Fig. 1; Table 1). Thus, under harsh high-inoculum conditions, no single time exposure achieves a universal kill guarantee. Instead, the framework successfully quantifies the conservative upper bound on residual viability and makes this isolate-level variability explicit.

**Fig. 1.**
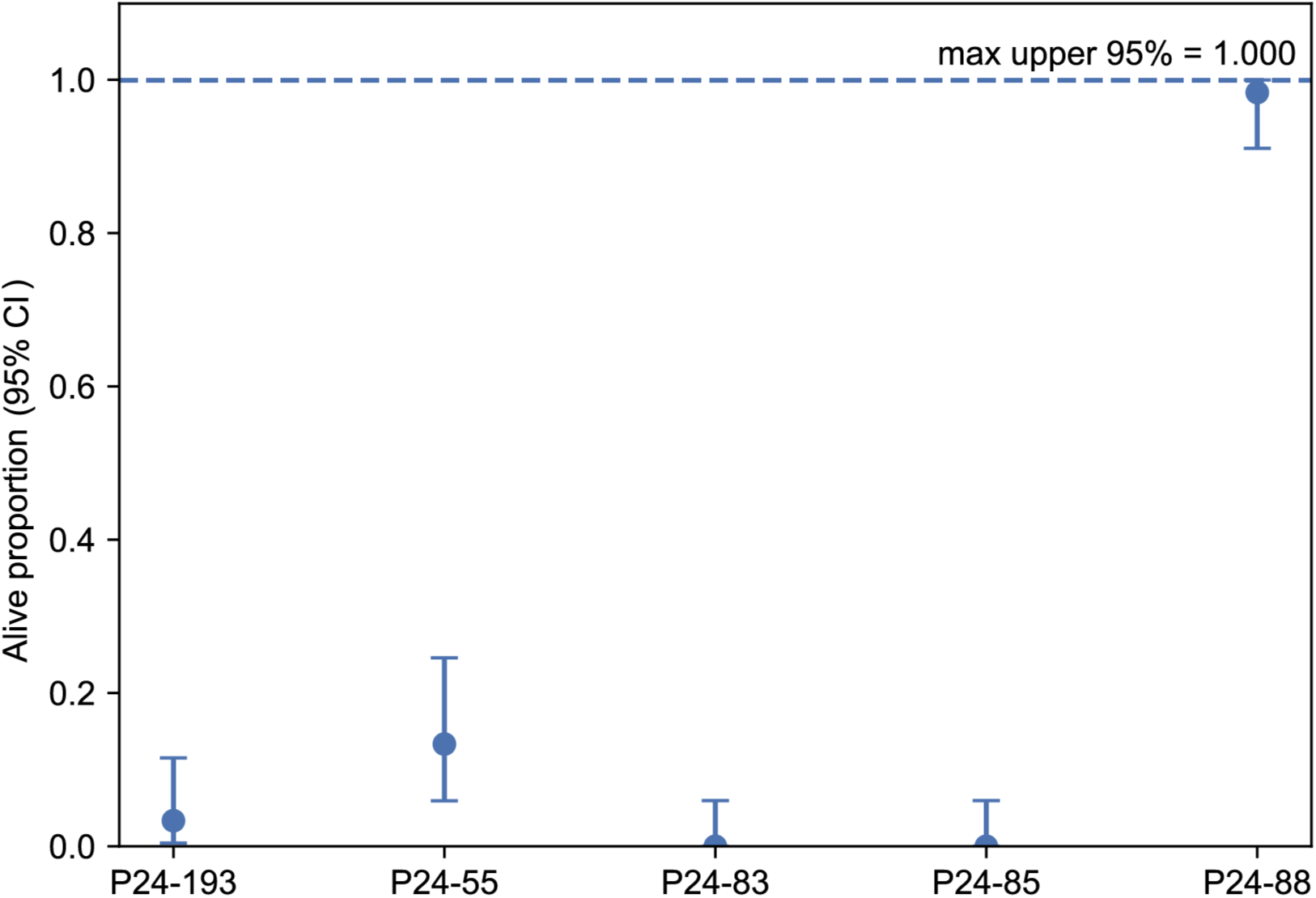
Isolate-specific alive proportion of Coffee *Colletotrichum* following an approximately 10 min UV-C exposure. Alive proportion for coffee *Colletotrichum* isolates following ∼10 min UV-C exposure. Dots represent the observed proportion (*p*-hat) of ‘alive’ colonies (survival ratio ≥ 0.05). Error bars show the exact Clopper-Pearson 95% CIs. The maximum upper 95% bound reached 1.000 (for isolate P24-88), demonstrating that no universal kill guarantee is achieved at this exposure, and highlighting critical isolate-specific heterogeneity.

**Table 1.** Probabilistic survival counts and confidence intervals for the Coffee UV-C cohort. Underlying data for Fig. 1, detailing the probabilistic survival analysis of the Coffee *Colletotrichum* cohort after 10 min UV-C exposure. Columns show the isolate identifier, total colony count (n), number of ‘alive’ colonies (defined as survival ratio ≥ 0.05), the observed proportion 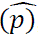, and the exact Clopper-Pearson 95% CI bounds.

| Isolate | n | Alive | $\hat{p}$ | CP 95 low | CP 95 up |
| --- | --- | --- | --- | --- | --- |
| P24-193 | 60 | 2 | 0.0333 | 0.00406 | 0.115 |
| P24-55 | 60 | 8 | 0.133 | 0.0594 | 0.246 |
| P24-83 | 60 | 0 | 0 | 0 | 0.0596 |
| P24-85 | 60 | 0 | 0 | 0 | 0.0596 |
| P24-88 | 60 | 59 | 0.983 | 0.911 | 1.0 |

Importantly, this probabilistic framework does not obscure biological reality. While many isolates were completely inactivated, qualitative inspection revealed significant isolate-specific heterogeneity. Notably, isolate P24-88 consistently exhibited localized survival as a dense central core or a thin translucent aggregation, whereas other isolates showed complete sterilization under the same UV-C regimen (Fig. 2). This demonstrates that our approach correctly quantifies both the overall efficacy and the critical isolate-level tail-risk that a single “average” metric would miss.

**Fig. 2.**
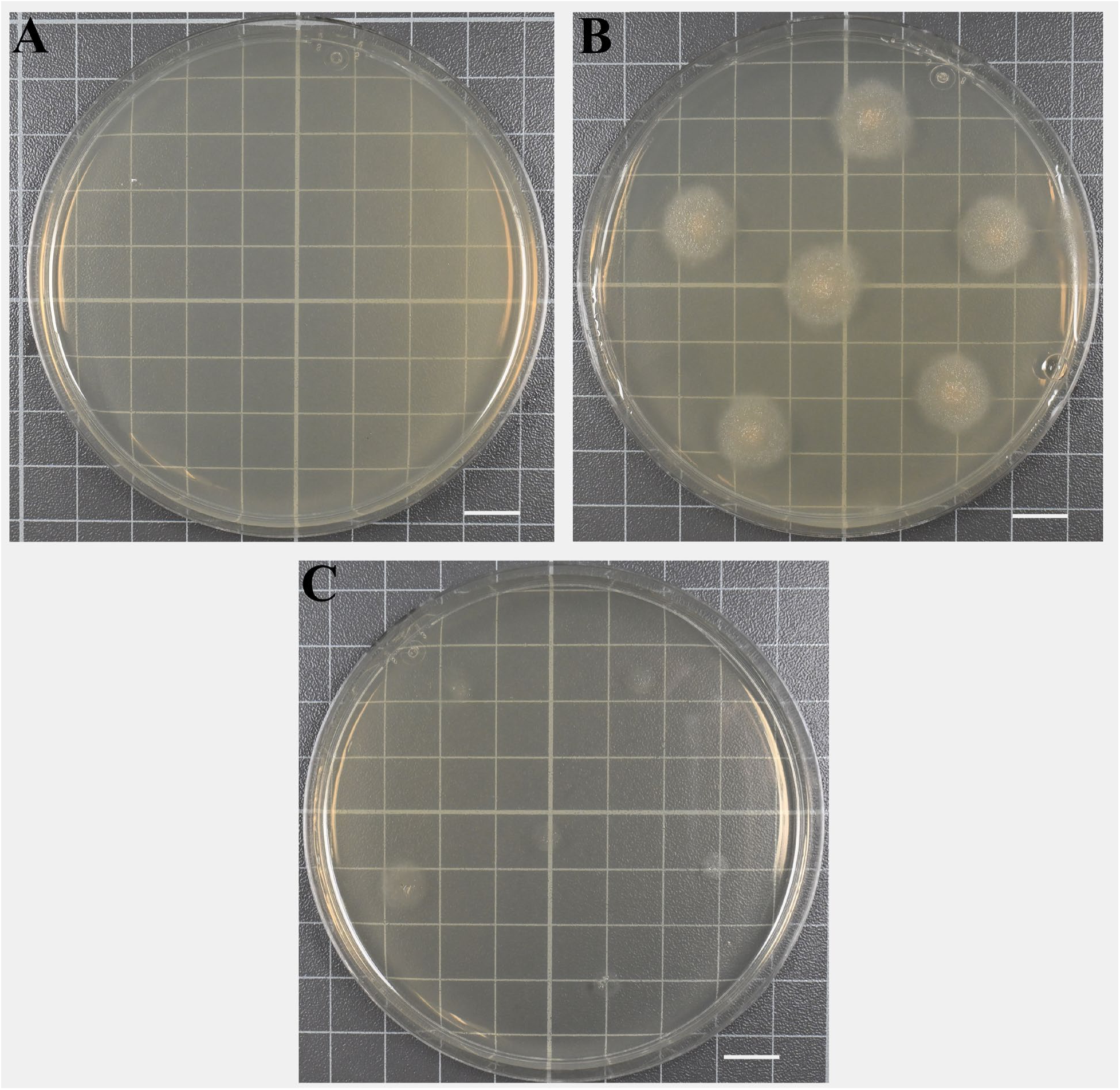
Representative plate images illustrating isolate-specific heterogeneity in the Coffee UV-C cohort, imaged at 48 hours post-inoculation. (A) Complete inactivation — six inoculation points are no longer visible (P24-55). (B) Untreated control showing normal colony growth (P24-88). (C) ∼10 min UV-C–treated plate showing localized survival P24-88) manifested as a dense central core or a thin translucent aggregation under identical exposure conditions. Scale bar = 1 cm.

At 275 nm and 0.58 mW cm⁻², the ∼10 min exposure corresponds to H = 348 mJ cm⁻² and a photon fluence of N = 4.82 × 10¹⁷ photons cm⁻², providing a reproducible baseline for translating “minutes” into energy and photons. Physics-informed lower bounds. Using the isolate-wise Clopper–Pearson upper 95 % bounds on survival at ∼10 min as S_up_and the single-hit inactivation model S = exp(−σN), we derived conservative lower bounds on the lethal cross-section 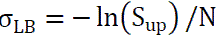. These σ_LB_ values span orders of magnitude across isolates, quantifying why a single time-point can never be universally protective (Fig. 3).

**Fig. 3.**
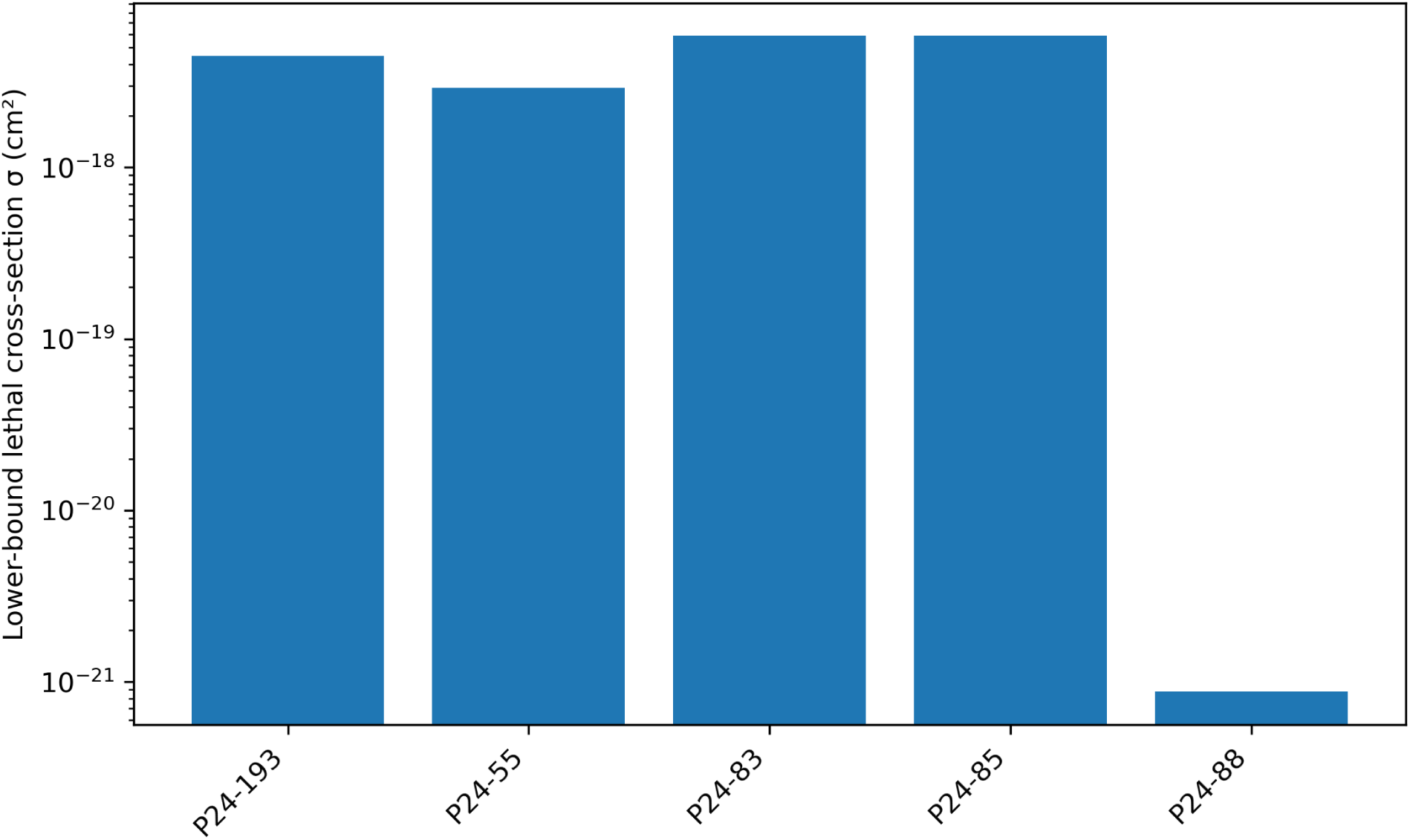
Conservative lethal cross-section lower bounds by isolate, derived from the Coffee cohort at ∼10 min UVC. Lower-bound 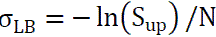 derived from isolate-wise Clopper–Pearson upper 95 % survival bounds at ∼10 min. (0.58 mW cm⁻²; 275 nm).

Risk-to-time translation. From σ_LB_, we computed conservative times required to reach 90/99/99.9 % inactivation at 0.58 mW cm⁻²; some isolates require minutes, whereas others are effectively unbounded under this irradiance (Fig. 4).

**Fig. 4.**
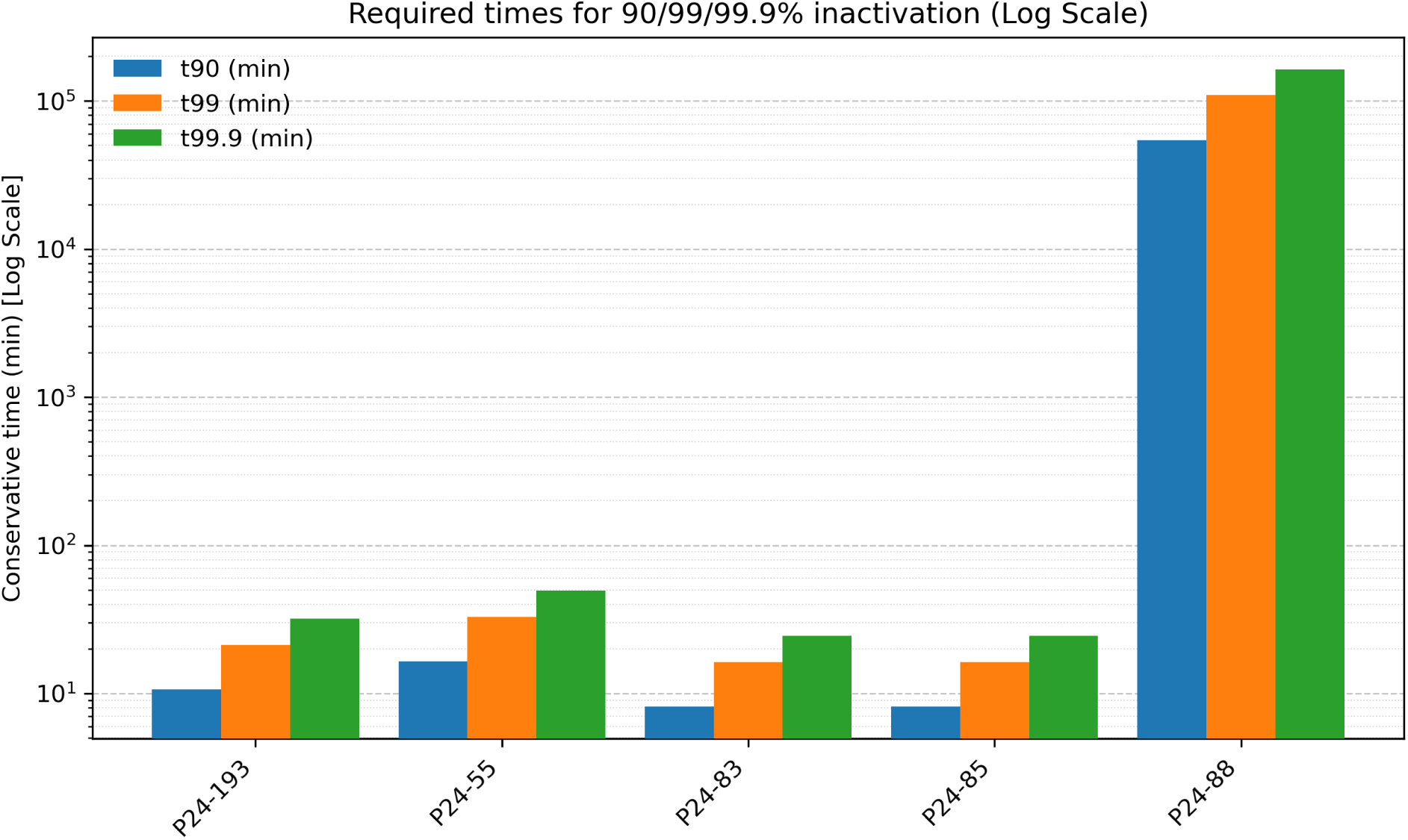
Required times for 90/99/99.9% inactivation at 0.58 mW cm⁻², plotted on a logarithmic Y-axis. Conservative inactivation times (*t*_*x*_) calculated from lower-bound lethal cross-sections (σ_*LB*_) for each isolate. The logarithmic scale is used to visualize the large variation across isolates with finite inactivation times. Note that infinite time values (e.g., t_99.9_ for P24-88, corresponding to S_up_) cannot be plotted on a log scale and are therefore omitted.

Burden penalty. Finally, an energy-per-expected-kill vs. inoculum density curve shows a steep penalty at higher spore loads; isolates with large S_up_ (e.g., P24-88) incur 2–3 orders-of-magnitude higher energy per kill than more susceptible isolates at the same exposure (Fig. 5).

**Fig. 5.**
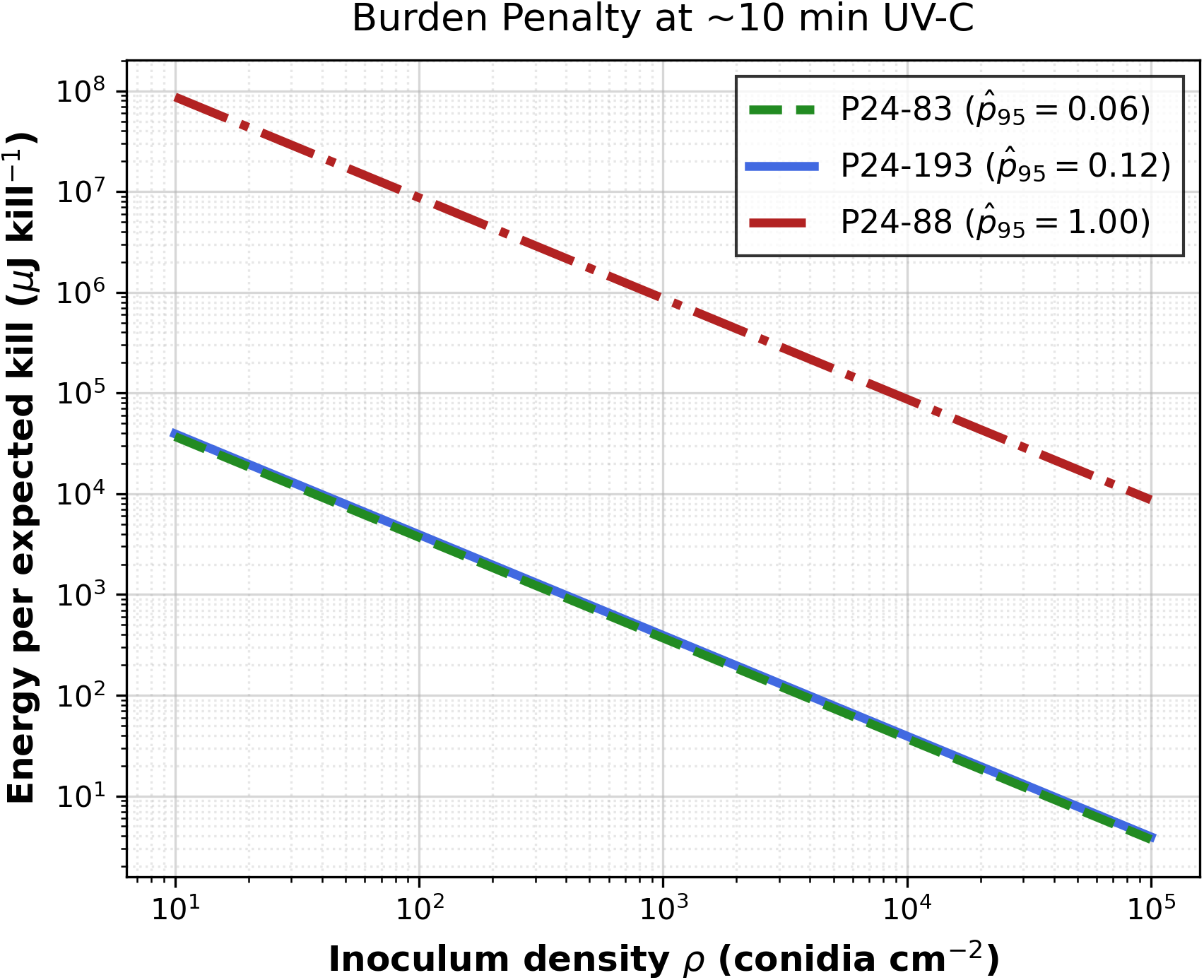
Energy per expected kill vs. inoculum density (µJ/kill) at ∼10 min. Upper-bound E/kill 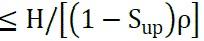 for representative isolates; note the steep burden penalty and the elevated curve for P24-88 (S_up_ = 1.00).

### 3.2. A size-free morphological fingerprint reveals nested pathosystem structure

Our second objective was to determine if fungal morphology encodes a “fingerprint” of stress that is independent of simple size reduction and generalizable across different stressors. For this analysis, we used the full dataset (n = 5,363), intentionally including the Cacao panel’s diverse UV-C, UV-B, and sonication treatments.

We performed PCA using only the three shape features (LWR, circularity, and IS&CG), explicitly excluding all size metrics (area, length, etc.). The results show that the Cacao and Coffee pathosystems occupy nested, rather than distinct, regions within this “shape space.” The Coffee cohort’s morphotypes are almost entirely contained within the much broader morphological space occupied by the Cacao cohort (Fig. 6). This demonstrates an asymmetric relationship—a host-specific geometric signature suggesting the Coffee pathosystem may represent a specialized subset derived from the wider geometric diversity present in the Cacao system.

**Fig. 6.**
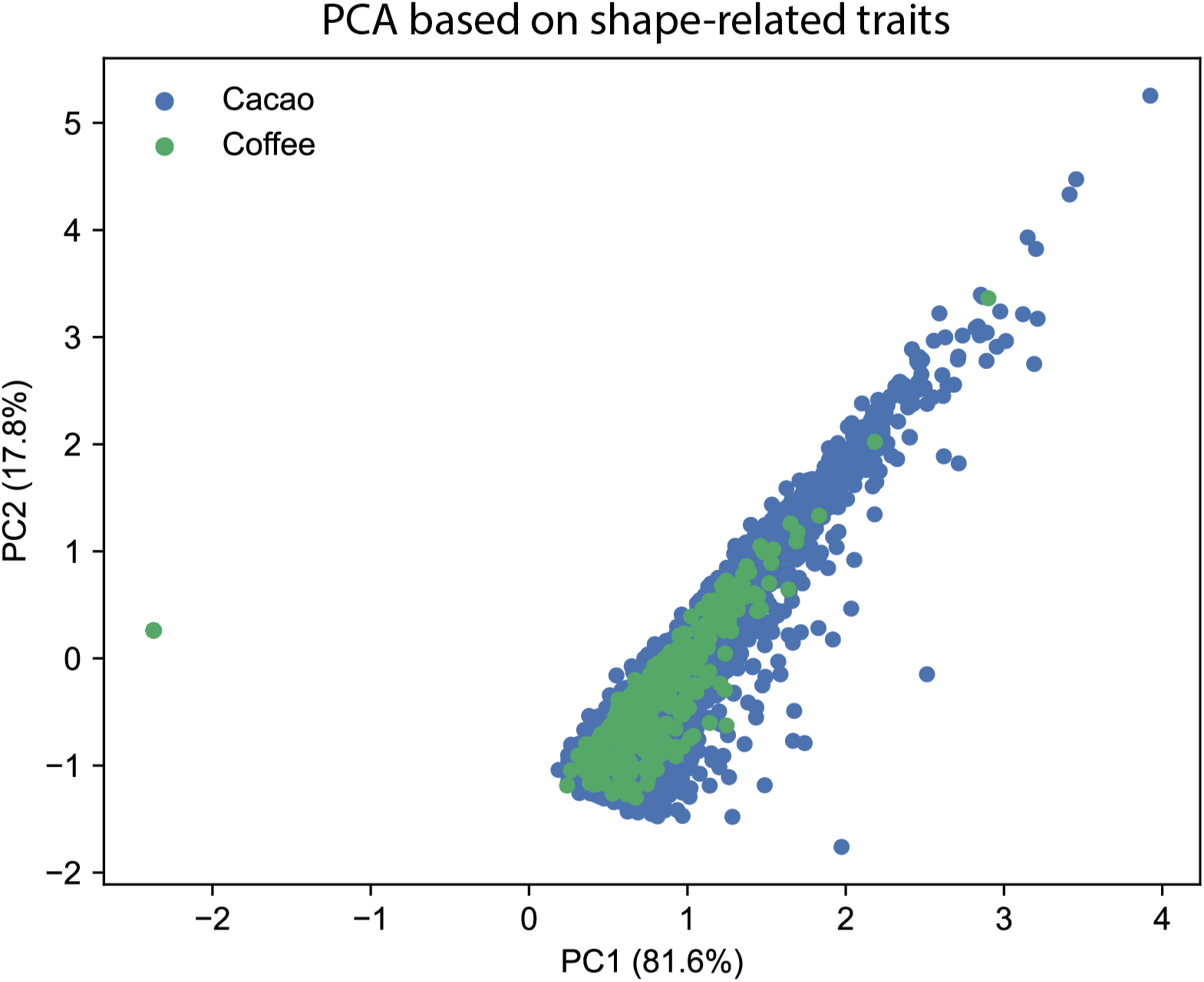
A size-free morphological fingerprint reveals nested relationships between pathosystems. PCA based only on shape features (LWR, circularity, and IS&CG). PCA colored by pathosystem, showing that the Coffee cohort’s morphospace (green) is nested within the broader Cacao cohort’s space (blue).

Univariate ANOVA confirmed that these shape features, particularly LWR and circularity, were highly significant drivers of treatment response across the entire dataset (Table 2). This confirms that geometry, not just growth, carries a transferable signal of stress response.

**Table 2.**
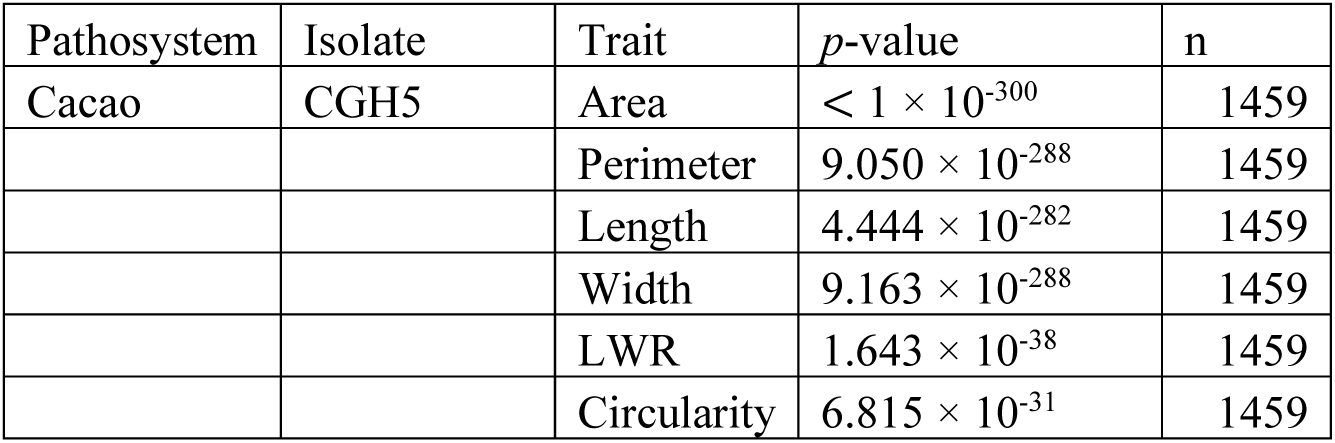
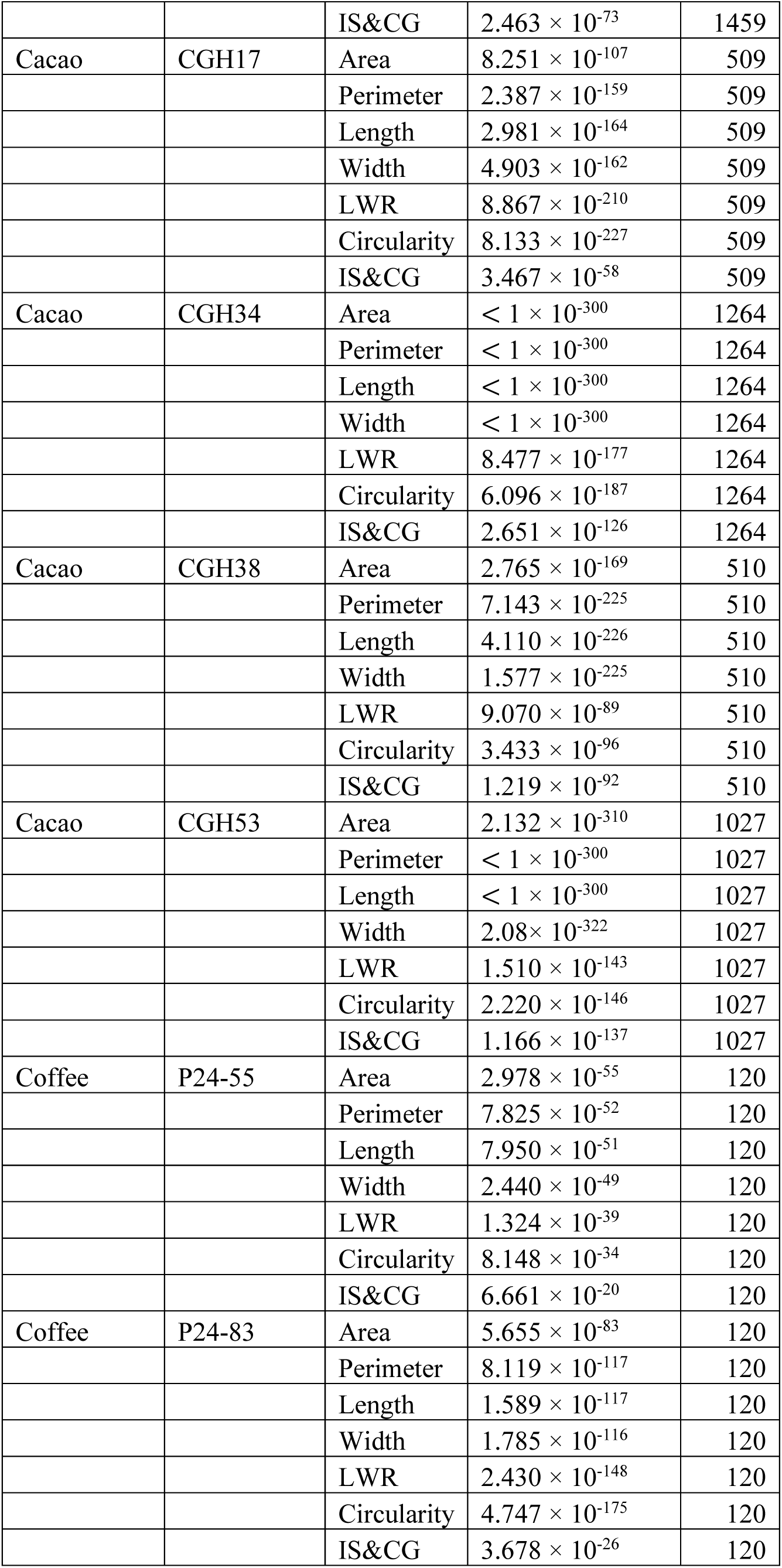
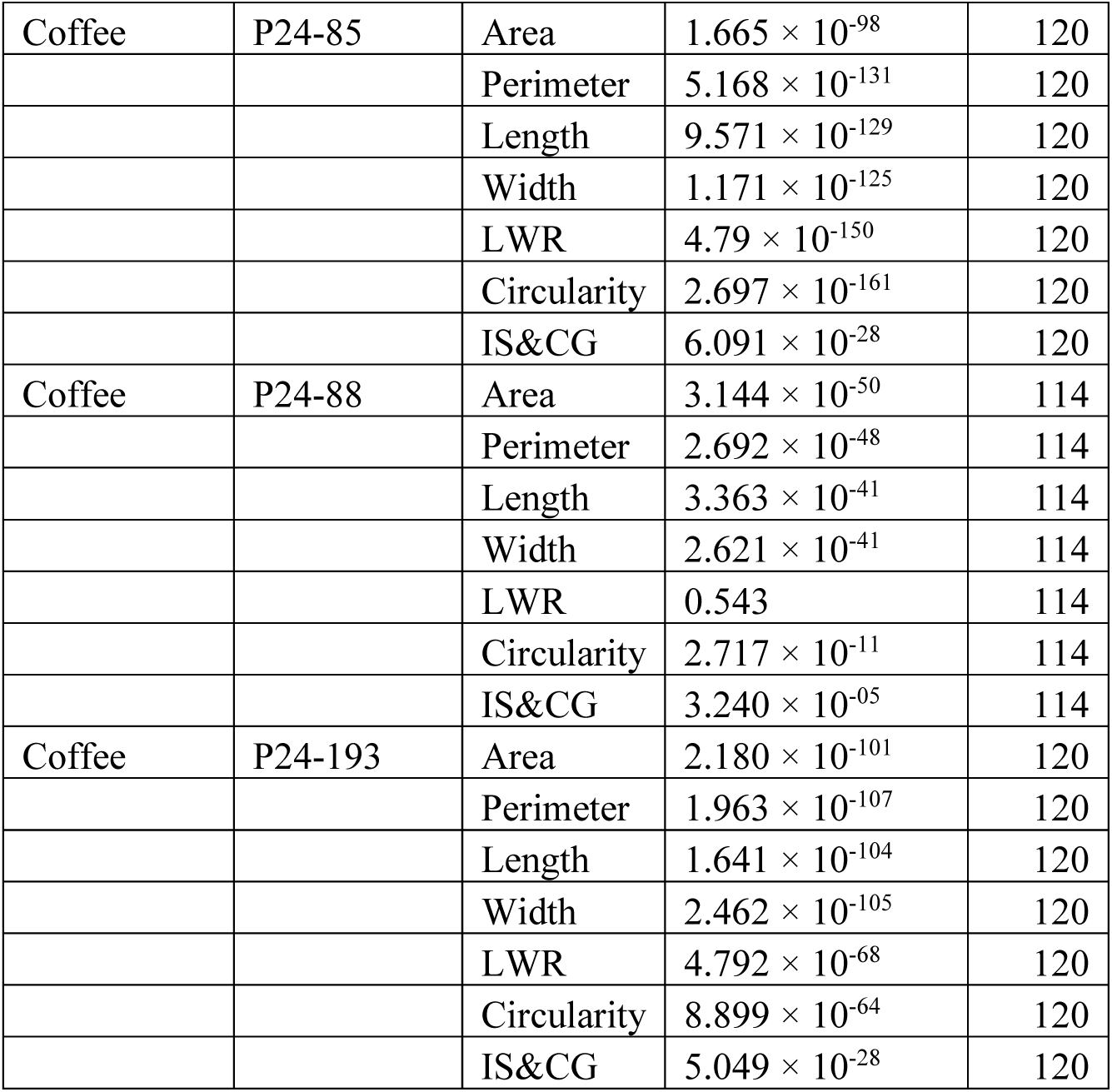
Univariate ANOVA results for treatment effects on morphological traits. Summary of one-way ANOVA (Type II SS) testing the effect of ‘Treatment’ (Control vs. All Treated conditions, including UV-C, UV-B, and Sonication) on each morphometric trait, performed independently for each isolate. The key finding is that “shape-only” features (LWR, circularity, and IS&CG) are also independently and highly significant, confirming that geometry, not just growth reduction, carries a transferable stress signal independent of pathosystem.

| Pathosystem | Isolate | Trait | *p*-value | n |
| --- | --- | --- | --- | --- |
| Cacao | CGH5 | Area | $< 1 \times 10^{-300}$ | 1459 |
| | | Perimeter | $9.050 \times 10^{-288}$ | 1459 |
| | | Length | $4.444 \times 10^{-282}$ | 1459 |
| | | Width | $9.163 \times 10^{-288}$ | 1459 |
| | | LWR | $1.643 \times 10^{-38}$ | 1459 |
| | | Circularity | $6.815 \times 10^{-31}$ | 1459 |
| | | IS&CG | $2.463 \times 10^{-73}$ | 1459 |
| Cacao | CGH17 | Area | $8.251 \times 10^{-107}$ | 509 |
| | | Perimeter | $2.387 \times 10^{-159}$ | 509 |
| | | Length | $2.981 \times 10^{-164}$ | 509 |
| | | Width | $4.903 \times 10^{-162}$ | 509 |
| | | LWR | $8.867 \times 10^{-210}$ | 509 |
| | | Circularity | $8.133 \times 10^{-227}$ | 509 |
| | | IS&CG | $3.467 \times 10^{-58}$ | 509 |
| Cacao | CGH34 | Area | $< 1 \times 10^{-300}$ | 1264 |
| | | Perimeter | $< 1 \times 10^{-300}$ | 1264 |
| | | Length | $< 1 \times 10^{-300}$ | 1264 |
| | | Width | $< 1 \times 10^{-300}$ | 1264 |
| | | LWR | $8.477 \times 10^{-177}$ | 1264 |
| | | Circularity | $6.096 \times 10^{-187}$ | 1264 |
| | | IS&CG | $2.651 \times 10^{-126}$ | 1264 |
| Cacao | CGH38 | Area | $2.765 \times 10^{-169}$ | 510 |
| | | Perimeter | $7.143 \times 10^{-225}$ | 510 |
| | | Length | $4.110 \times 10^{-226}$ | 510 |
| | | Width | $1.577 \times 10^{-225}$ | 510 |
| | | LWR | $9.070 \times 10^{-89}$ | 510 |
| | | Circularity | $3.433 \times 10^{-96}$ | 510 |
| | | IS&CG | $1.219 \times 10^{-92}$ | 510 |
| Cacao | CGH53 | Area | $2.132 \times 10^{-310}$ | 1027 |
| | | Perimeter | $< 1 \times 10^{-300}$ | 1027 |
| | | Length | $< 1 \times 10^{-300}$ | 1027 |
| | | Width | $2.08 \times 10^{-322}$ | 1027 |
| | | LWR | $1.510 \times 10^{-143}$ | 1027 |
| | | Circularity | $2.220 \times 10^{-146}$ | 1027 |
| | | IS&CG | $1.166 \times 10^{-137}$ | 1027 |
| Coffee | P24-55 | Area | $2.978 \times 10^{-55}$ | 120 |
| | | Perimeter | $7.825 \times 10^{-52}$ | 120 |
| | | Length | $7.950 \times 10^{-51}$ | 120 |
| | | Width | $2.440 \times 10^{-49}$ | 120 |
| | | LWR | $1.324 \times 10^{-39}$ | 120 |
| | | Circularity | $8.148 \times 10^{-34}$ | 120 |
| | | IS&CG | $6.661 \times 10^{-20}$ | 120 |
| Coffee | P24-83 | Area | $5.655 \times 10^{-83}$ | 120 |
| | | Perimeter | $8.119 \times 10^{-117}$ | 120 |
| | | Length | $1.589 \times 10^{-117}$ | 120 |
| | | Width | $1.785 \times 10^{-116}$ | 120 |
| | | LWR | $2.430 \times 10^{-148}$ | 120 |
| | | Circularity | $4.747 \times 10^{-175}$ | 120 |
| | | IS&CG | $3.678 \times 10^{-26}$ | 120 |
| Coffee | P24-85 | Area | $1.665 \times 10^{-98}$ | 120 |
| | | Perimeter | $5.168 \times 10^{-131}$ | 120 |
| | | Length | $9.571 \times 10^{-129}$ | 120 |
| | | Width | $1.171 \times 10^{-125}$ | 120 |
| | | LWR | $4.79 \times 10^{-150}$ | 120 |
| | | Circularity | $2.697 \times 10^{-161}$ | 120 |
| | | IS&CG | $6.091 \times 10^{-28}$ | 120 |
| Coffee | P24-88 | Area | $3.144 \times 10^{-50}$ | 114 |
| | | Perimeter | $2.692 \times 10^{-48}$ | 114 |
| | | Length | $3.363 \times 10^{-41}$ | 114 |
| | | Width | $2.621 \times 10^{-41}$ | 114 |
|  |  | LWR | 0.543 | 114 |
| | | Circularity | $2.717 \times 10^{-11}$ | 114 |
| | | IS&CG | $3.240 \times 10^{-05}$ | 114 |
| Coffee | P24-193 | Area | $2.180 \times 10^{-101}$ | 120 |
| | | Perimeter | $1.963 \times 10^{-107}$ | 120 |
| | | Length | $1.641 \times 10^{-104}$ | 120 |
| | | Width | $2.462 \times 10^{-105}$ | 120 |
| | | LWR | $4.792 \times 10^{-68}$ | 120 |
| | | Circularity | $8.899 \times 10^{-64}$ | 120 |
| | | IS&CG | $5.049 \times 10^{-28}$ | 120 |

### 3.3. Shape-only features predict both host-origin and survival outcomes

To quantify the predictive strength of this size-free fingerprint, we trained two machine-learning models on the full dataset (n = 5,363) using only the shape features and time/fluence variables, with strict plate-grouped cross-validation.

***Host-origin classification***: A Gradient Boosting Classifier distinguished cacao vs. coffee isolates with Accuracy = 0.933 ± 0.010 and a Macro-F1 = 0.819 ± 0.012. This confirms that the geometric fingerprint is not only statistically significant but robustly predictive of the pathogen’s origin system, even without size cues.

***Survival-ratio regression***: A Gradient Boosting Regressor trained on the same features predicted the continuous survival ratio for all treated plates (UV-B/C, sonication) with R² = 0.742 ± 0.009 and RMSE = 0.204 ± 0.014. The model thus provides a quantitative decision support for anticipating residual viability across treatments. Together, these results demonstrate that shape alone, independent of size, conveys a generalizable signal capable of predicting both a pathogen’s identity and its post-stress fate.

## 4. Discussion

Our findings fundamentally challenge the conventional use of fixed-time UV-C recipes (e.g., “10 minutes”) for postharvest control. Such heuristics are dangerously simplistic as they fail to account for the vast, isolate-specific heterogeneity endemic to fungal pathogens. We demonstrated that a ∼10 min exposure on the Coffee cohort did not yield a universal kill. Instead, it revealed a conservative upper 95% confidence bound on survival of 1.000 for isolate P24-88 (Fig. 1), quantifying the extreme risk posed by this heterogeneity. This study replaces time-based heuristics with a dual framework that is both predictive and diagnostic: (1) a physics-informed probabilistic model that translates “minutes” into device-agnostic energy quanta, quantifying risk from heterogeneity and inoculum density, and (2) a machine-learning-based morphological fingerprint that validates the physiological state.

The translation of “10 minutes” into a reproducible physical dose (H = 348 mJ cm⁻²; N = 4.82 × 10¹⁷ photons cm⁻²) is the first step toward a device-agnostic protocol. Our decision to model inactivation kinetics using a single-hit model (S = e^−σN^) is not arbitrary; this first-order kinetic approach is a well-established photobiological standard for the UV inactivation of fungal conidia (Norman, 1951). This model allowed us to calculate a conservative lower-bound lethal cross-section (σ_*LB*_), a fundamental parameter representing the effective “target size” of the pathogen (Fig. 3). The results were stark: σ_*LB*_ values spanned orders of magnitude. For P24-88, the high survival (S_up_ ≈ 1.0) yielded a σ_LB_ so low that the calculated time to achieve 99.9% inactivation (t_99.9_) effectively diverges to infinity under this irradiance (Fig. 4). This saturation effect mirrors findings in clinical disinfection, where increasing UV-C fluence beyond a threshold (e.g., >300 mJ cm⁻²) often fails to yield additional log reductions due to organic interference or shadowing (Wallace et al., 2019). Similarly, in static juice treatments, insufficient mixing leads to survival tails, whereas dynamic flows (e.g., Dean vortex) significantly enhance inactivation (Müller et al., 2011), supporting our finding that static shading is a critical barrier.

This “tailing” phenomenon, where a resistant sub-population survives extreme doses, is a known challenge in disinfection (Gardner and Shama, 1998). Our findings provide a clear biological hypothesis for this resistance. P24-88’s resilience is likely due to efficient photoprotection, consistent with high concentrations of pigments like 1,8-dihydroxynaphthalene (DHN)-melanin, which are known to confer significant UV-C tolerance in fungi (Yang et al., 2022). This isolate-specific heterogeneity is not an anomaly; it is the central challenge in *Colletotrichum* control. This same heterogeneity in response to physical (UV) (Baek et al., 2025b), chemical (novel phenolic amides) (Ahn et al., 2025), and biological (*Trichoderma*) stressors (Baek et al., 2025a) are reported. The extreme UV-C resistance of P24-88 is, therefore, a critical data point in a much larger pattern of pathogen-specific adaptation. These observed ‘tails’ in survival probability likely reflect the pathogen’s inherent physiological plasticity, a biological barrier that simple time-based treatments fail to address. Our probabilistic approach effectively encapsulates this biological uncertainty into a safety margin, ensuring that control measures remain robust even against evolved resilience.

Furthermore, our model provides a physical explanation for the localized survival of P24-88 in dense conidial cores. The E/kill model (Fig. 5) demonstrates a steep “burden penalty”: as inoculum density (ρ) increases, the energy required per kill rises exponentially. This is consistent with the well-documented “shielding” effect, where spores in aggregates or dense masses physically block UV radiation, protecting the cells underneath (Gardner and Shama, 1998; Rotem and Aust, 1991). The survival rate of aggregated spores has been shown to be directly proportional to the number of conidia in the mass (Rotem and Aust, 1991). This finding provides a critical, actionable lever for practical control: managing the initial inoculum burden (e.g., via pre-rinsing) may be as important as increasing the UV dose or exposure time.

From an operational standpoint, our results argue that UV-C should not be deployed as a stand-alone, one-size-fits-all recipe, but as a calibrated component of integrated postharvest systems. The physics-informed risk maps (Figs. 3–5) highlight that managing inoculum burden, for example through pre-washing or biological antagonists, can be as impactful as increasing UV-C dose for reducing tail-risk, especially in heavily contaminated lots. Furthermore, the shape-only fingerprint provides a rapid, non-destructive diagnostic for monitoring fungal stress and residual viability after treatment. Together, these elements connect the biological basis of UV-C tolerance to actionable control levers, aligning with broader efforts to cut postharvest food loss and ensure the safety of horticultural products.

Our second framework, morphological fingerprinting, addresses the challenge of identifying this stress state. UV exposure is not merely a binary kill/no-kill event but also induces morpho-developmental changes; for instance, UV radiation is known to affect appressorium formation and microcycle conidiation (Ghajar et al., 2006). This justifies the analysis of spore morphology as a proxy for stress response. Automated image analysis workflows are increasingly recognized for their ability to quantify size and shape distributions that manual microscopy misses, revealing environmental influences on spore maturation (Woyzichovski et al., 2021). Our Gradient Boosting models, using only size-invariant shape features, successfully classified both the host-of-origin (Cacao vs. Coffee) and, more importantly, the survival outcome (live vs. dead) across diverse stressors (UV-C, UV-B, Sonication).

This approach is validated by a growing body of research applying machine learning to fungal classification. Using CNNs and other deep learning models to classify fungi based on microscopic images of spores and morphology is an established, state-of-the-art method (Prommakhot and Srinonchat, 2024; Tahir et al., 2018).

Our laboratory has previously applied this core conceptual approach, using machine learning to analyze stress-induced shape fingerprints, to decode an ‘ecological memory’ of host-origin from chemical stress phenotypes (Baek et al., 2025c). This work demonstrates that the exact shape fingerprint is also robust in predicting the outcome of physical stress (UV-C), confirming its utility as a universal, stress-agnostic diagnostic tool. Integrating such morphometric data with molecular phylogenetics and multi-omics is envisioned as the future roadmap for resolving complex fungal taxonomies and resistance traits (Naqvi et al., 2025).

While all experiments were conducted under high-inoculum in vitro conditions, the probabilistic framework is designed to be calibratable. Future work will focus on translating these confidence bounds to fruit surfaces, accounting for wound architecture and senescence status. Although focused here on cacao and coffee *Colletotrichum*, this shape-based, probabilistic approach is directly portable to other postharvest pathosystems (e.g., mango anthracnose or citrus green mold), provided that their UV response and morphotypes are quantified in a comparable manner.

In conclusion, this dual framework provides a comprehensive solution. The physics-informed risk model (Figs. 3-5) moves postharvest control from simple heuristics to quantifiable, risk-based protocols that account for the critical variables of isolate heterogeneity (e.g., melanin content) and inoculum density. The ML-based morphological fingerprint provides a rapid, validated method to assess fungal stress response, demonstrating that shape is a major indicator of pathogen physiology along with size.

## 5. Conclusion

We bifurcated the problem of UV-C control:

1. For practitioners: We replaced ambiguous “minutes” with auditable confidence bounds. In the Coffee ∼10 min cohort, the upper 95% survival bound reached 1.000, demonstrating no universal guarantee at that setting and motivating risk-aware adjustments (fluence, duration, adjuncts) tailored to heterogeneity.
2. For scientists/QC: We showed that a size-free, shape-only fingerprint generalizes across stressors and is predictive of both host origin (Accuracy ≈ 0.933; Macro-F1 ≈ 0.819) and survival (R² ≈ 0.742), enabling device-agnostic phenotyping and monitoring.

Together, these contributions deliver a transparent, extensible platform that (i) turns postharvest UV-C prescriptions from minutes into probabilistic bounds, and (ii) elevates geometry from a descriptive by-product to a decision-grade signal for managing isolate-level variation in fungal control.

## CRediT authorship contribution statement

Ezekiel Ahn: Writing – original draft, Supervision, Project administration, Methodology, Formal analysis, Data curation, Conceptualization. Insuck Baek: Writing – review & editing, Validation, Software, Resources. Seunghyun Lim: Writing – review & editing, Formal analysis. Amelia Lovelace: Writing – review & editing, Resources. Minhyeok Cha: Writing – review & editing, Resources. Moon S. Kim: Writing – review & editing, Supervision, Resources, Funding acquisition. Sunchung Park: Writing – review & editing, Methodology. Lyndel W. Meinhardt: Writing – review & editing, Supervision, Resources, Funding acquisition.

## Author agreement statement

Author Agreement Statement We the undersigned declare that this manuscript is original, has not been published before and is not currently being considered for publication elsewhere. We confirm that the manuscript has been read and approved by all named authors and that there are no other persons who satisfied the criteria for authorship but are not listed. We further confirm that the order of authors listed in the manuscript has been approved by all of us. We understand that the Corresponding Author is the sole contact for the Editorial process.

## Declaration of Generative AI and AI-assisted technologies in the writing process

During the preparation of this work, the authors used Google Gemini in order to improve the readability and language quality of the manuscript. After using this tool/service, the author(s) reviewed and edited the content as needed and take full responsibility for the content of the publication.

## Acknowledgments

Mention of any trade names or commercial products in this article is solely for the purpose of providing specific information and does not imply recommendation or endorsement by the U. S. Department of Agriculture. USDA is an equal opportunity provider and employer, and all agency services are available without discrimination. This work is supported by the U.S. Department of Agriculture, Agricultural Research Service, In-House Projects No. 8042-21220-258-000-D and 8042-21000-303-000-D.

## Appendix A. Supplementary material

The raw and processed datasets supporting the conclusions of this article are provided in the Supplementary Information (Supplementary Data 1).

Supplementary Data 1. Comprehensive dataset and analysis outputs. This file contains the complete numerical results supporting the study, organized into the following sheets:

S1–S2 (Morphometrics): Descriptive statistics (mean, SEM) and ANOVA results for morphological traits across all isolates and treatments.

S3 (Model Applicability): Clarification on the single-dose experimental design and the consequent exclusion of multi-parameter curve fitting.

S4 (Raw Data): The integrated isolate-level dataset (n = 5,363), combining the newly generated Coffee UV-C cohort (∼10 min) with the Cacao reference panel for cross-pathosystem modeling.

S5–S8 (Physics & Probability): Calculated Clopper–Pearson 95% confidence intervals (CP95) for survival and corresponding lower-bound lethal cross-sections ($\sigma_{LB}$) for the Coffee UV-C cohort.

S9–S10 (Machine Learning): Performance metrics (Accuracy, F1-score, R², RMSE) for host-origin classification and survival regression models.

S11 (Parameters): Physical constants and irradiance parameters used for photon fluence calculations.

### Data availability

The complete Python script used to perform all analyses, including data preprocessing, physics modeling, and machine learning pipelines, will be made publicly available in the project’s GitHub repository upon acceptance of the manuscript.

